# Structural basis for Fc receptor recognition of immunoglobulin M

**DOI:** 10.1101/2022.11.18.517032

**Authors:** Qu Chen, Rajesh Menon, Pavel Tolar, Peter B. Rosenthal

## Abstract

FcμR is the IgM-specific Fc receptor involved in the survival and activation of B cells. Using cryo-EM, we reveal eight binding sites for the human FcμR Ig domain on the IgM pentamer, one of which overlaps with the receptor binding site for the transcytosis receptor pIgR, but a different mode of binding explains Ig isotype specificity. The complex explains engagement with polymeric serum IgM and the monomeric IgM B cell receptor.

## Main text

Immunoglobulin Fc receptors are effector molecules expressed on the surface of immune cells, which can generate a wide range of protective functions crucial in immune responses after engaging with the Fc domains of the immunoglobulins. FcμR (historically also called TOSO or FAIM3) is a high-affinity Fc receptor specific for IgM^1^. Although IgM is a primordial Ig isotype present in all vertebrates, FcμR has a relatively late appearance during early mammalian evolution^2^. Human FcμR is composed of an extracellular immunoglobulin (Ig)-like domain, a flexible stalk, a transmembrane region and a long cytoplasmic tail^3^. The Ig-like domain is responsible for ligand binding^3^ and shares about 40% sequence identity with the first Ig-like domain of polymeric immunoglobulin receptor (pIgR-D1), which is encoded by a gene located in the same chromosomal region in mammals^2^. However, rather than binding to both polymeric IgM and IgA as pIgR does^4–7^, FcμR exclusively binds to IgM^8^, indicating its specific binding mechanisms as well as functional roles. Potential residues responsible for the binding between IgM and FcμR have been proposed but the structure of the FcμR/IgM complex is currently unknown^9–11^.

Here we study a complex of the ectodomain of human FcμR with the IgM-Fc pentameric core. The IgM core closely resembles previously described pentameric IgM structures^4,6,12^, consisting of Cμ3 and Cμ4 domains, assembled at the C-terminal tailpieces with the J chain (Fig. 1a). Four FcμR binding sites are observed at the front of subunits Fcμ1 to Fcμ4 (Fig. 1a) and four at the back of subunits Fcμ2 to Fcμ5 (Fig. 1b, data processing workflow in Extended Data Fig. 1-2). The other two potential binding sites at the front of Fcμ5 and the back of Fcμ1 are likely to be blocked due to the proximity of the hairpin-1 and hairpin-2 loops of the J chain, respectively. Only the Ig-like domain of FcμR is ordered, indicating flexibility of the stalk region mediating membrane attachment. Each FcμR is similarly positioned relative to the IgM subunit (Extended Fig. 3), however with different occupancies (Fig. 1a-b, Extended Fig. 4, Extended Data Table 1) and adaption to asymmetric features of the IgM structure. Only Cμ4 domains of IgM are involved in the interactions. Superposition of subunits of the IgM-FcμR complex on the IgM-BCR ^13–15^, which contains a monomeric IgM (mIgM) identical to the subunits of IgM pentamer at Cμ4 domains, and two signalling chains Ig*αβ*, shows that mIgM-BCR can accommodate binding of two FcμR Ig domains (Fig. 1c).

**Fig. 1.**
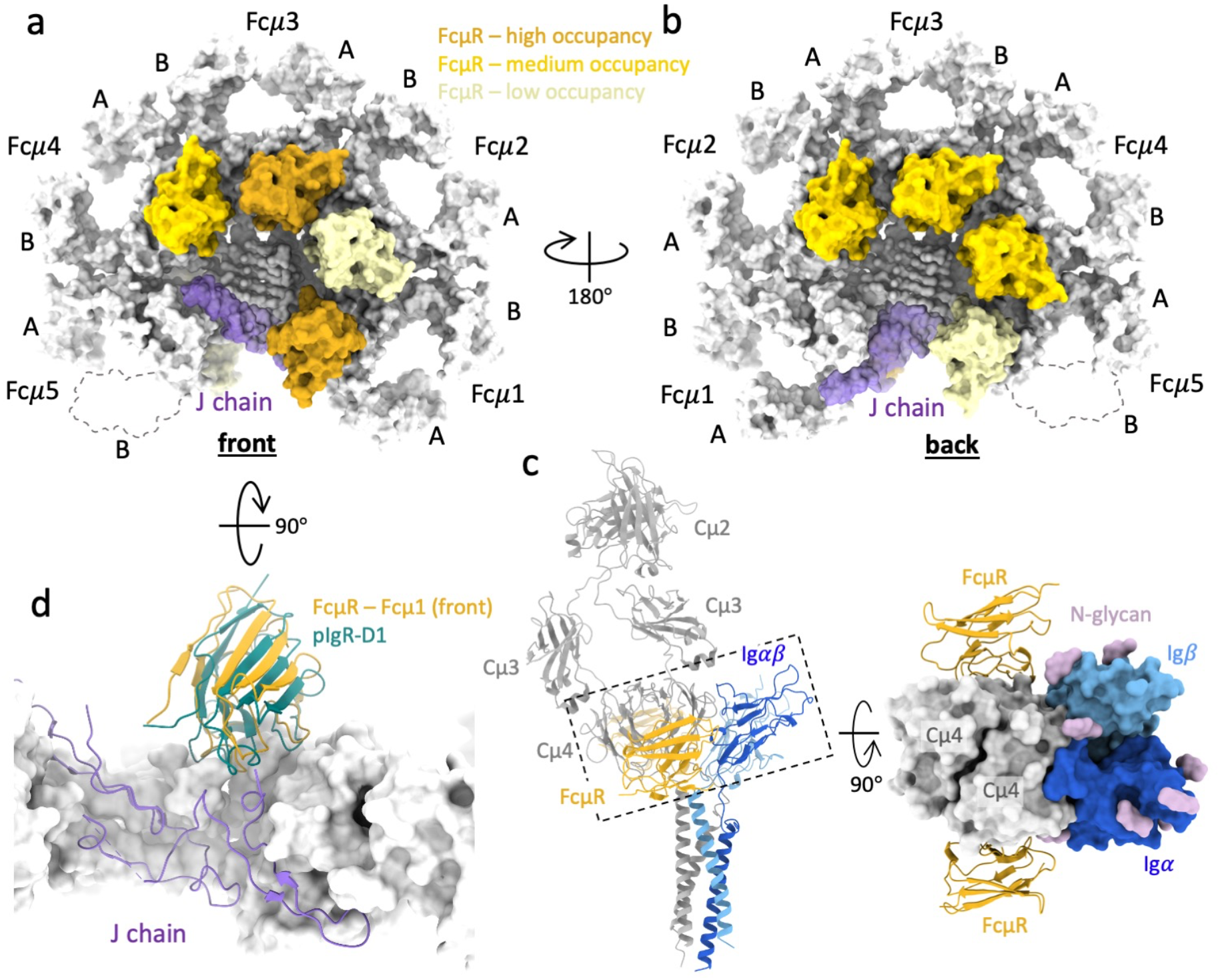
Multivalent binding of FcμR to pentameric IgM. (a-b) Structure of the human FcμR/IgM-Fc complex showing four FcμR binding at each side (a, front; b, back) of the IgM-Fc platform. Cμ3, Cμ4 and tailpieces of IgM in grey, J chain in purple. FcμR in dark, medium, and pale yellow indicate high, medium, and low occupancies of FcμR at different binding sites. The Cμ3-B domain in subunit Fcμ5, which is removed by proteolysis, is shown in dashed contour. (c) Overlay of the IgM-BCR (pdb id 7XQ8) and a subunit of IgM pentamer with two FcμR bound at both sides, aligned at the Cμ4 domains. Right panel, the top view of the region highlighted in the dashed box in the left panel shows no steric clash or interactions between FcμR and Ig*αβ* (in blue) on either side. (d) Overlapped binding sites for pIgR-D1 (cyan) and FcμR at subunit Fcμ1, aligned at IgM, revealing a lifted position of FcμR compared to pIgR-D1.

Two binding sites (front of subunit Fcμ1 and Fcμ3, darker yellow in Fig. 1a), which have highest occupancies of FcμR, reached high-resolution (3.5 Å and 3.1 Å) in the cryo-EM maps (Extended Fig. 5 and 6), thus allowing the atomic model interpretation of the FcμR Ig-like domain and the IgM binding interface. The Ig-like domain of FcμR contains several conserved structural features with pIgR-D1, including two intrachain disulfide bonds (Cys49-Cys58 and Cys37-Cys104), a salt bridge between Arg75 and Asp98, as well as three loops analogous to the complementarity-determining regions (CDR) of immunoglobulin variable domains, which are responsible for engaging IgM (Fig. 2a).

**Fig. 2.**
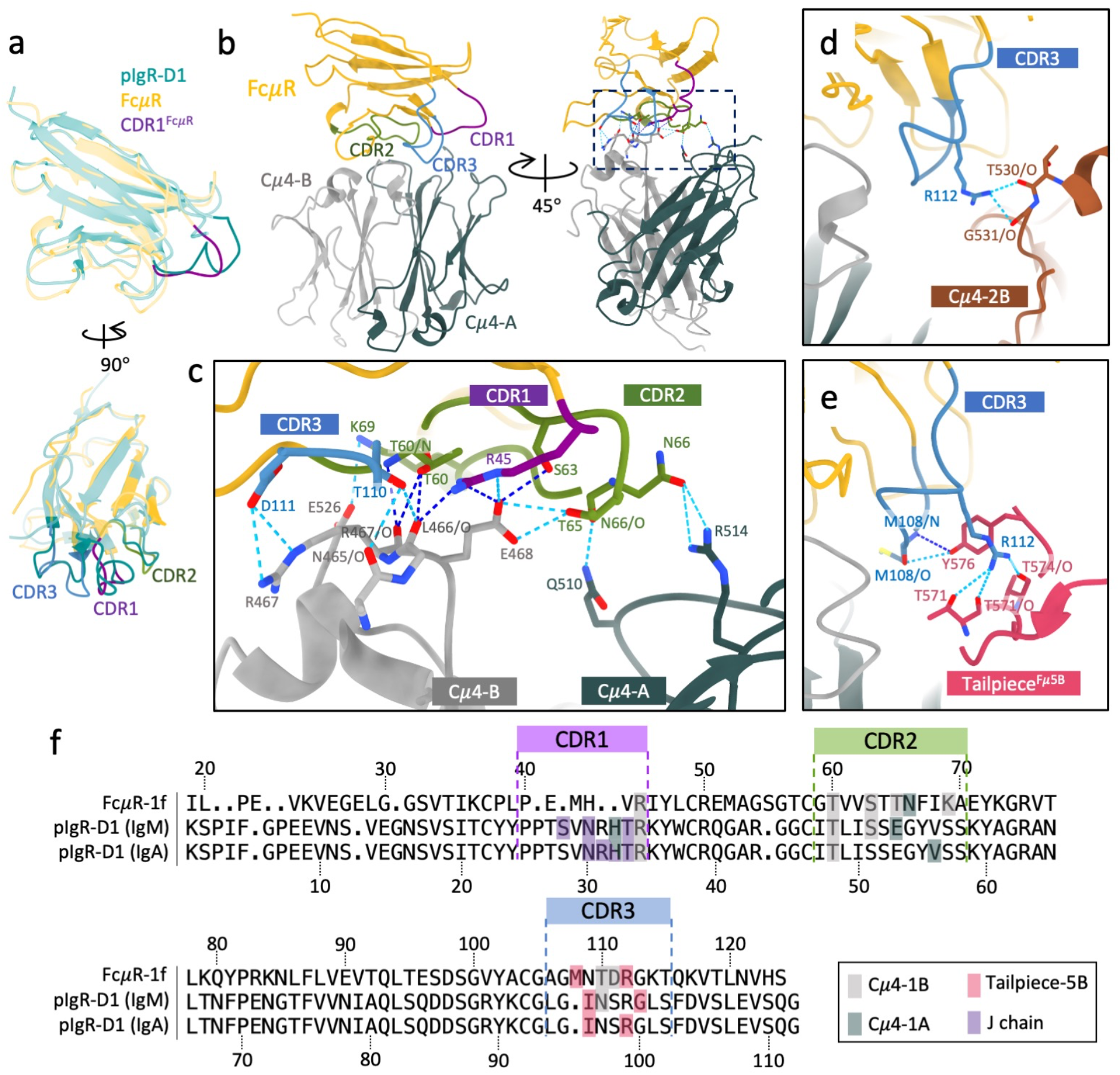
The binding interface between FcμR and IgM. (a) Superimposed FcμR and pIgR-D1 with the three CDR loops highlighted in FcμR. CDR1, purple; CDR2, green; CDR3, blue. (b) The interactions between the three CDR loops of FcμR and the two Cμ4 domains of the subunit Fcμ3. Cμ4-A in slate grey, Cμ4-B in light grey. (c) Zoom-in view of the region highlighted in the dashed box in (b) showing the interacting residues on the three CDR loops of FcμR and the two Cμ4 domains. The interactions displayed are at subunit Fcμ3 but also shared in subunit Fcμ1. The hydrogen bonds (H-bonds) are represented by dashed lines in blue. Dark blue lines are conserved H-bonds between pIgR-D1 and IgM, and the light blues are unique in FcμR/IgM interface. (d) Interacting of CDR3 of the FcμR at subunit Fcμ3 with the neighbouring Cμ4 domain in IgM subunit Fcμ2 (in brown). (e) Interacting of CDR3 of the FcμR at subunit Fcμ1 with the tailpiece in the B chain of subunit Fcμ5 (in pink). (f) Structural-based sequence alignment of FcμR at subunit Fcμ1 (first row) and pIgR-D1 (second row) highlighting residues interacting with the indicated regions of IgM. The third row shows pIgR-D1 interactions with IgA.

The FcμR at the front of Fcμ1 overlaps with the pIgR-D1 binding sites on IgM, but in a slightly lifted position relative to the IgM (Fig. 1d) and 25% smaller buried surface area (Extended Data Table 2). This difference is likely due to the truncated CDR1 region in FcμR (Fig. 2a). Only residue, Arg45, at the tip of CDR1 of FcμR, is conserved in pIgR-D1 (Arg34), whereas the other key CDR1 residues in pIgR-D1 which interact intensively with the J-chain are missing in FcμR. The CDR2 and CDR3 regions of the two receptors are structurally similar and interact with similar regions on IgM (Fig. 2a-c, f).

Most of the interacting residues on IgM are shared between the FcμR at subunit Fcμ1 and Fcμ3, illustrated in Fig. 2c (Extended Fig. 7). Residues Asn465-Glu468 on Cμ4-B chain of the IgM subunit form a central hub for FcμR binding, which interacts with all three CDR loops on FcμR (Fig. 2c). Cμ4-A has two residues (Gln510 and Arg514) interacting with CDR2 on FcμR (Fig. 2c). Glu510 was identified as a binding site for FcμR and IgM by mutagenesis^10^. The main differences between the FcμR binding sites at subunit Fcμ1 and Fcμ3 are at CDR3. At subunit Fcμ3, Arg112 in FcμR interacts with two carbonyl groups (Thr530 and Gly531) on the Cμ4-2B domain in the neighbouring IgM subunit (Fig. 2d), which is presumably shared by all the FcμR bound at internal subunits (subunit Fcμ2-4). The particularly high occupancy of FcμR at subunit Fcμ3 is likely due to the additional contact from the carbohydrate chain extending from Asn563 on the tailpiece of subunit Fcμ3 (Extended Data Fig. 8). Nevertheless, it has been found that FcμR can still engage with de-glycosylated IgM and trigger internalisation by the cells^9^, suggesting redundancy thanks to the multivalent binding of FcμR to IgM. In contrast, the FcμR at the front of subunit Fcμ1 is stabilised by multiple interactions with the tailpiece on subunit Fcμ5 extended from the other side of the IgM core (Fig. 2e), which is also observed in the binding interface between pIgR-D1 and IgM but with different residues (Fig. 2f).

The interactions observed between the CDR loops on FcμR and the IgM core provide a structural basis for the IgM specificity of FcμR. With truncated CDR1 region, FcμR binding on IgM is predominantly stabilised by CDR2 and CDR3. On the other hand, CDR1 contributes significantly to pIgR binding to both Ig isotypes (Fig. 2f), and greater for IgA than IgM based on the percentage of buried surface area involved (35.2% vs 26.5%, Extended table 2). Fewer residues on pIgR-D1 in CDR2 and CDR3 regions are involved in interactions with IgA than IgM (four vs six, Fig. 2f). As a result, a truncated CDR1 would have a greater impact on receptor binding to IgA than IgM. In addition, two of the residues on IgM responsible for FcμR binding (Arg514 and Arg467) are not conserved in IgA (Glu363 and Asn362), which would potentially disrupt the interactions at Asn66 (CDR2), Lys69 (CDR2) and Asp111 (CDR3) in FcμR. These differences between the two Ig isotypes likely account for FcμR specificity for IgM.

We showed the structural basis for multiple FcμR binding sites on IgM. Multivalent engagement of IgM facilitates capture of soluble IgM or IgM immune complexes by FcμR anchored on cell surfaces, leading to clustering, signalling^16^, and transport^17^ through FcμR. Clustering has been found to be essential to induce phosphorylation at the serine and tyrosine residues in the immunoglobulin tail tyrosine (ITT) motif within the cytoplasmic region of FcμR^1^. The complex structure also explains the observed colocalisation of FcμR and IgM-BCR on membranes of mature B cells that promote B cell survival^18^ and on the trans-Golgi network (TGN) in developing B cells which regulates the transport of IgM-BCR from TGN to the B cell surface^19^. A key common feature of the FcμR binding sites is that only the Cμ4 domains are involved in the interactions. They therefore recognise a structurally invariant region, whereas other parts of IgM, including Cμ3, may vary in structure^12,14^. Overall, our structures reveal the binding and isotype specificity of FcμR for IgM and promote the understanding of the functions of FcμR in the BCR signalling pathway.

## Methods

### pIgM-Fc/Fc*μ*R complex sample preparation

Full-length IgM (myeloma, Jackson Immunoresearch, 009-000-012) was HPLC purified using Superose 10/300 column with the running buffer (Tris-HCl buffer 50 mM, 11.5 mM CaCl^2,^ pH 8.1). The IgM sample was then concentrated to around 2 mg/mL and ultrapure trypsin with equivalent 4% weight of IgM was added to the sample. The digestion reaction was kept at 56°C for 30 min. The digested sample was then run again on HPLC column again with PBS as running buffer to isolate the correct IgM-Fc fragment, and fractions was double-checked by visualising on 3-8% Tris Acetate protein gels. The IgM-Fc sample was diluted to 1 mg/ml in PBS. Lyophilized human Fc*μ*R (R&D, Catalog 9494-MU-050) was suspended in PBS to 1 mg/ml. The binding of IgM to surface-immobilised Fc*μ*R was validated by biolayer interferometry with sub-nanomolar affinity. pIgM-Fc and Fc*μ*R were mixed with equal volume, which corresponds to molar ratio about 1:10. The mixture was kept at room temperature for about 10 min.

### pIgM-Fc/Fc*μ*R complex cryo-EM grid preparation

Quantifoil (300 Cu mesh, R2/2) was washed by chloroform, dried in air, and glow discharged with air (25 mA, 30 s). The mixture was then diluted 10-fold and 4 ul of diluted sample was immediately pipetted to the glowed grid in the environmental chamber of a Vitrobot Mark IV (FEI/Thermo) at 4 °C and 100% humidity. The grid was blotted for 4 s before plunged into liquid ethane kept at liquid nitrogen temperature.

#### Cryo-EM data collection

The IgM cryo grids were firstly screened on Talos Arctica microscope (FEI/Thermo) at 200 kV and best ones were transferred to a Titan Krios microscope (FEI/Thermo) at 300 kV equipped with a Gatan Imaging Filter (GIF) using EPU software (v 2.11). The slit width of the energy filter was set to 20 eV. 17,835 movies were recorded on a K2 camera in counting mode with a total dose of 50.6 electrons per Å^2^ fractionated over 40 frames (dose rate 5.06 e^-^/Å^2^/s) with a 1.08 Å pixel size and a defocus range between -1.2 to -3.5 μm. 16,713 more movies with the same conditions above and with 20°stage tilting were then collected to help to improve the particle orientation distribution.

#### Cryo-EM data processing

The workflow of the Cryo-EM data processing is shown in Extended Data Fig. 1a. Both non-tilted and tilted movies were imported into Relion (v 3.1)^20^, followed by Relion’s own motion correction and CTF estimation (CTFFIND, v 4.1.13)^21^. For tilted dataset, patchCTF in CryoSPARC (v 3.2.0)^22^ was also conducted in parallel on the aligned micrographs to obtain a better estimation of the defocus values. 5.9M particles in total were picked by a trained model in CrYOLO (v 1.7.5)^23^ from both dataset and extracted in Relion with box size of 100 pixels (bin4, pixel size=4.32 Å). The original CTF values of the particles in the tilted dataset was then substituted by the results from patchCTF by using the patchCTF extraction function in CryoSparc, and the particles were re-extracted in Relion with the same box size and binning as the non-tilted particles. The particles were then combined and subjected to 2D classification in CryoSPARC, with 2.08 M particles selected based on the populations and resolutions of the class averages. Typical 2D class averages are shown in Extended Data Fig. 1b. The selected particles were re-extracted again in Relion with box size of 400 pixels with original pixel size (1.08 Å). The particles were then refined, Bayesian polished, and refined again in Relion (unsharpened map in Extended Data Fig. 1c). The half-map FSC at 0.143 is 3.5 Å (FSC plot in Extended Data Fig. 1d).

To address problems caused by the quasi-two-fold symmetry of IgM pentamer and different occupancies of FcμR binding, particle subtraction was conducted to remove the signal of FcμR from the IgM core (pink mask in Extended Data Fig. 2a), followed by 3D classification without image alignment to separate the particle subset with best resolution at the IgM core. Highly resolved J chain is an indicator of good particle alignment. 516,875 particles from the 3D class highlighted in the green box in Extended Data Fig. 2a were selected and reverted to the original particles with all the signals restored. The particles were then 3D refined by non-uniform refinement in Cryosparc as well as CTF refinement options including beam tilt and per-particle defocus correction. The refined map reveals eight FcμR binding sites on IgM core with different intensities of FcμR, which sequentially appears when lowering the threshold of the Gaussian-filtered map (Extended Data Fig. 4a).

The relative intensities of densities at individual FcμR binding sites reflect the occupancy of FcμR at each position, which were quantified by focused 3D classification at individual subunits described in Extended Data Fig. 4b. Masks were created for individual subunits for particle subtraction, where only signal inside the masks remained. 3D classification without image alignment was then calculated for each subunit. The percentages of the binding states for each subunit (bound at front, back, both, or neither) are presented in Extended Data Fig. 4b and also summarised in Extended Data Table 1. It is worth noting that using different number of classes (from six to fifteen) for the 3D classification only had limited influence on the results, indicating that results are reliable.

To resolve the binding interfaces in atomic detail, focused 3D classifications and refinements were performed at the subunits with highest occupancies of FcμR, i.e., at subunit Fcμ1 and Fcμ3. The 3D classifications started from all the particles (2,084,147) in Extended Data Fig. 1 to preserve as many good particles as possible for further refinement. For subunit Fcμ3, two FcμR molecules are bound at both front and back of the subunit, resulting in a local C2 symmetry, which were implemented in 3D classification and refinement. The workflows of focused classification and refinement for subunit Fcμ1 and Fcμ3 are shown in Extended Data Fig. 5 and Extended Data Fig. 6, respectively. CTF refinement parameters including beam-tilt, and per-particle-defocus corrections are also applied to the non-uniform refinements.

#### Model building and refinement

The initial Atomic coordinate model of Fc*μ*R was predicted by trRosetta^24^, and the IgM-Fc cryo-EM model (pdb id 6KXS) was used as the initial model for IgM core.

For the map refined using the particles selected by the focused 3D classification at subunit Fc*μ*1 (Extended Data Fig. 4a), all ten Fcμ chains as well as the J chain in the IgM cryo-EM model (pdb id 6KXS) were used as the initial model. One FcμR was built in the density at the front of subunit Fcμ1. Real space refinement in Phenix (v 1.19.2)^25^ was performed before manual fixing the clashes and outliers in Coot (v 0.9.6)^26^. Iterations between auto-and manual-refinements were conducted for optimisation. N-Acetylglucosamine (NAG) molecules on the four IgM Fcμ chains at Asn563 were built and refined in Coot.

For the map refined using the particles selected by the focused 3D classification at subunit Fc*μ*3 (Extended Data Fig. 6a), only four Fcμ chains (chain C, D, E, and F) in the IgM-Fc cryo-EM model (pdb id 6KXS) were kept instead of the whole molecule, and two predicted FcμR models were added at the front and the back of IgM. The refinements were also performed in Phenix and Coot as described above.

Eight Fc*μ*R models were built into the map refined using the particles selected by the focused 3D classification at the IgM core (Extended Data Fig. 2a). The two models above were combined and the FcμR model refined at subunit Fcμ3 (models in yellow, Extended Data Fig. 6b) was also duplicated and fitted into the FcμR densities at the other five FcμR positions in subunit Fcμ2, Fcμ4, and Fcμ5. Rigid-body refinement was then performed in Phenix for the recombined model (IgM core + 8 FcμR).

All three cryo-EM maps were sharpened with corresponding models in LocScale^27^ in ccpem (v 1.6.0)^28^ with their corresponding models shown in Extended Data Fig. 2b, Extended Data Fig. 5b and Extended Data Fig. 6b.

### Map and model validation

A series of validations were conducted on the maps and models, shown in panel c to f in Extended Data Fig. 2, Extended Data Fig. 5, and Extended Data Fig. 6. The half-map Fourier shell correlation (FSC) for the three structures indicate 3.6 Å, 3.5 Å and 3.1 Å at 0.143 cut-off and map-model FSC plots show 3.9 Å, 3.6 Å and 3.3 Å at 0.5 cut-off (panel c). Local resolution and 3DFSC (panel d and e) were calculated in Cryosparc after refinement. 3DFSC results shown in Extended Data Fig. 2f and Extended Data Fig. 5f indicate some degree of angular anisotropy, caused by preferred orientation of the particles within the ice (Extended Data Fig. 2e and Extended Data Fig. 5e). Peptide chains were validated in Coot and Phenix and carbohydrates were validated using Privateer^29^ in ccpem. The data table for Cryo-EM data collection, processing, and validation statistics are summarised in Extended Data Table 3. The figures are made with UCSF Chimera (v 1.13.1)^30^ and UCSF ChimeraX (v 1.4)^31^.

## Data availability

The structural data that support the findings of this study have been deposited in the Protein Data Bank and EM Data bank. The 8:1 FcμR/IgM-Fc model displayed in Fig. 1 has entry number EMD-16150 and PDB-8BPE. The IgM-Fc core with one FcμR at subunit Fcμ1 has entry number EMD-16151 and PDB-8BPF. The IgM subunit Fcμ3 with two FcμR has entry number EMD-16152 and PDB-8BPG.

## Acknowledgements

We thank A. Nans of the Structural Biology Science Technology Platform for advice on data collection and computing; L. Masino, A. Purkiss and P. Walker of the Structural Biology Science Technology Platform; the Scientific Computing Science Technology Platform for computational support. This work was supported by the Francis Crick Institute, which receives its core funding from Cancer Research UK (FC001143 (P.B.R), FC001185 (P.T.)), the UK Medical Research Council (FC001143 (P.B.R), FC001185 (P.T.)), and the Wellcome Trust (FC001143 (P.B.R), FC001185 (P.T.)).

## Author Contributions

Q.C., R.M. performed experiments. Q.C., R.M., P.T., and P.B.R. contributed to experimental design, data analysis and manuscript writing.

## Competing interests

The authors declare no competing interests.

**Correspondence and requests** for materials should be addressed to P.T. and P.B.R.

---

**Extended Data Fig. 1.**
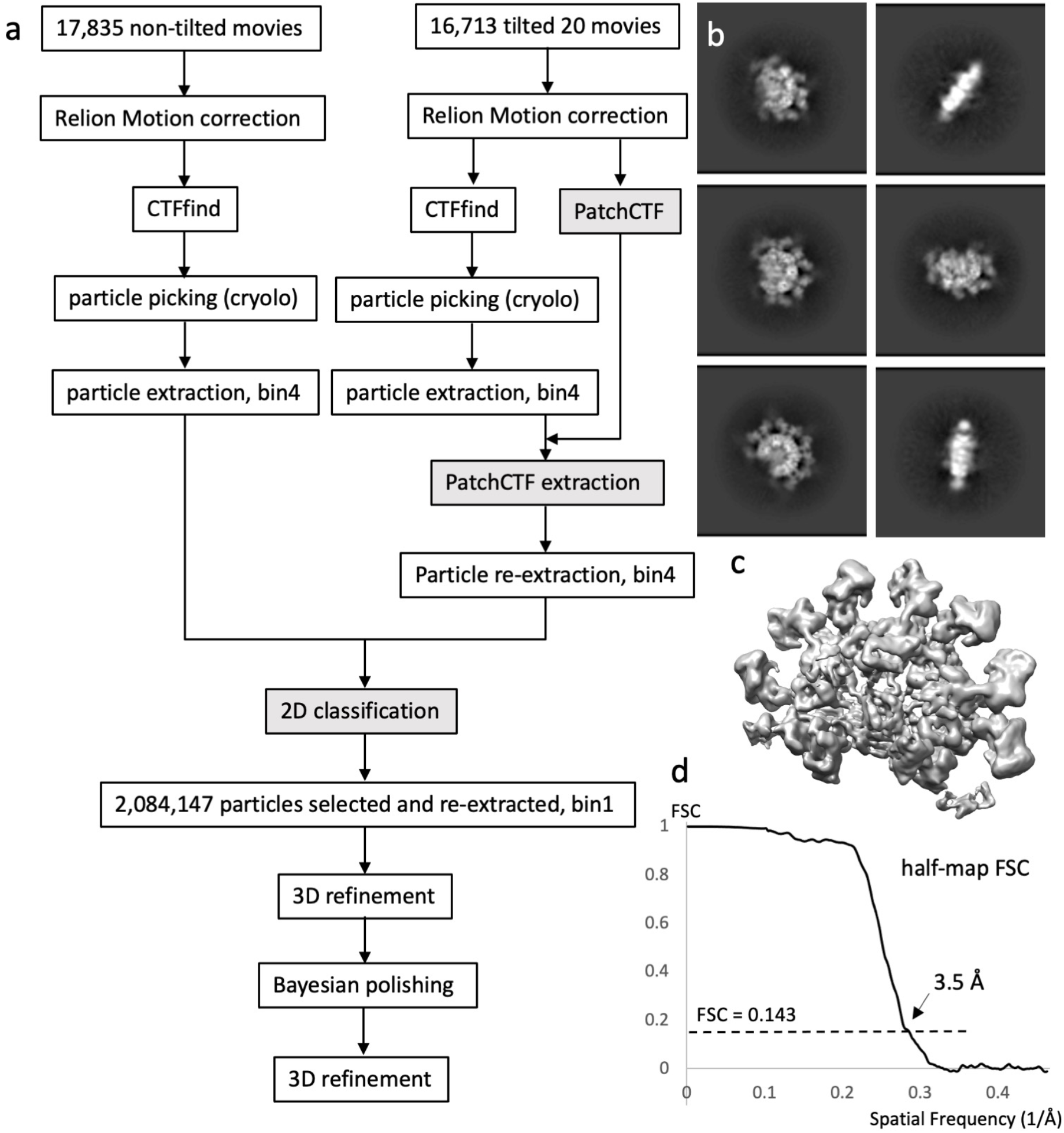
Single particle analysis of FcμR/IgM-Fc. (a) Flow chart of the data processing for both non-tilted and tilted datasets. Steps are conducted in Relion 3.1 (clear box) or Cryosparc 3.2.0 (grey box). (b) Typical 2D classes of the complex. (c) 3D auto-refined map. (d) Half-map Fourier shell correlation (FSC) plot showing 3.5 Å global resolution.

**Extended Data Fig. 2.**
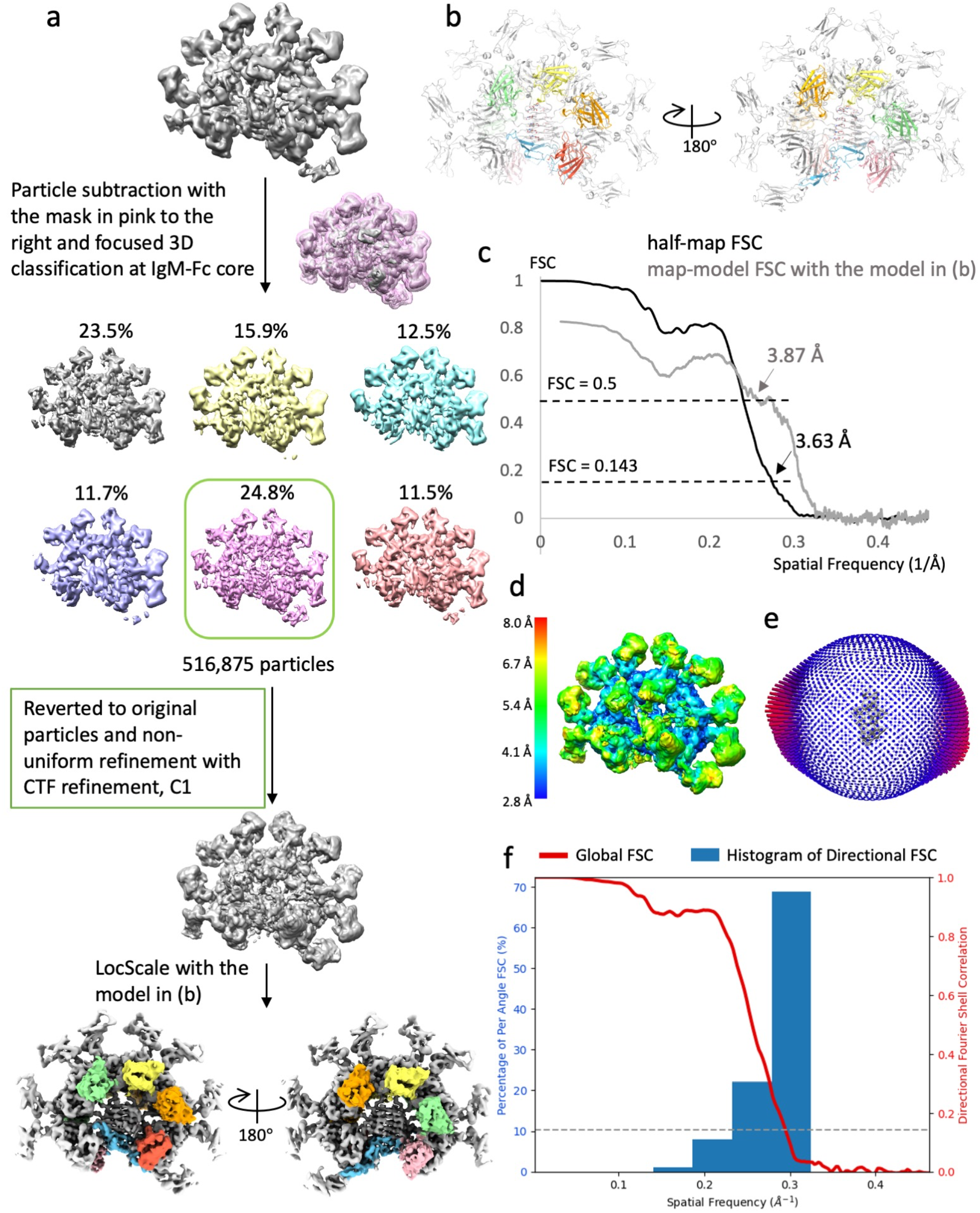
Cryo-EM structure of IgM-Fc/FcμR complex. (a) Particle subset selection by focused 3D classification at the IgM-Fc core and map refinement. (b) Front and side view of the complex model. IgM in grey, FcμR in dark blue. (c) Gold standard Fourier shell correlation (FSC) with 3.6 Å resolution at 0.143 cut-off and map-model FSC plot showing 3.9 Å resolution at 0.5 cut-off calculating with the model shown in (b) calculated in Phenix. (d) Local resolution of the refined map calculated in Cryosparc. (e) Eulerian angle distribution of the particles in the non-uniform refinement. (f) 3DFSC histogram calculated in Cryosparc of the refined map showing anisotropy between 3.2 Å - 5.4 Å.

**Extended Data Fig. 3.**
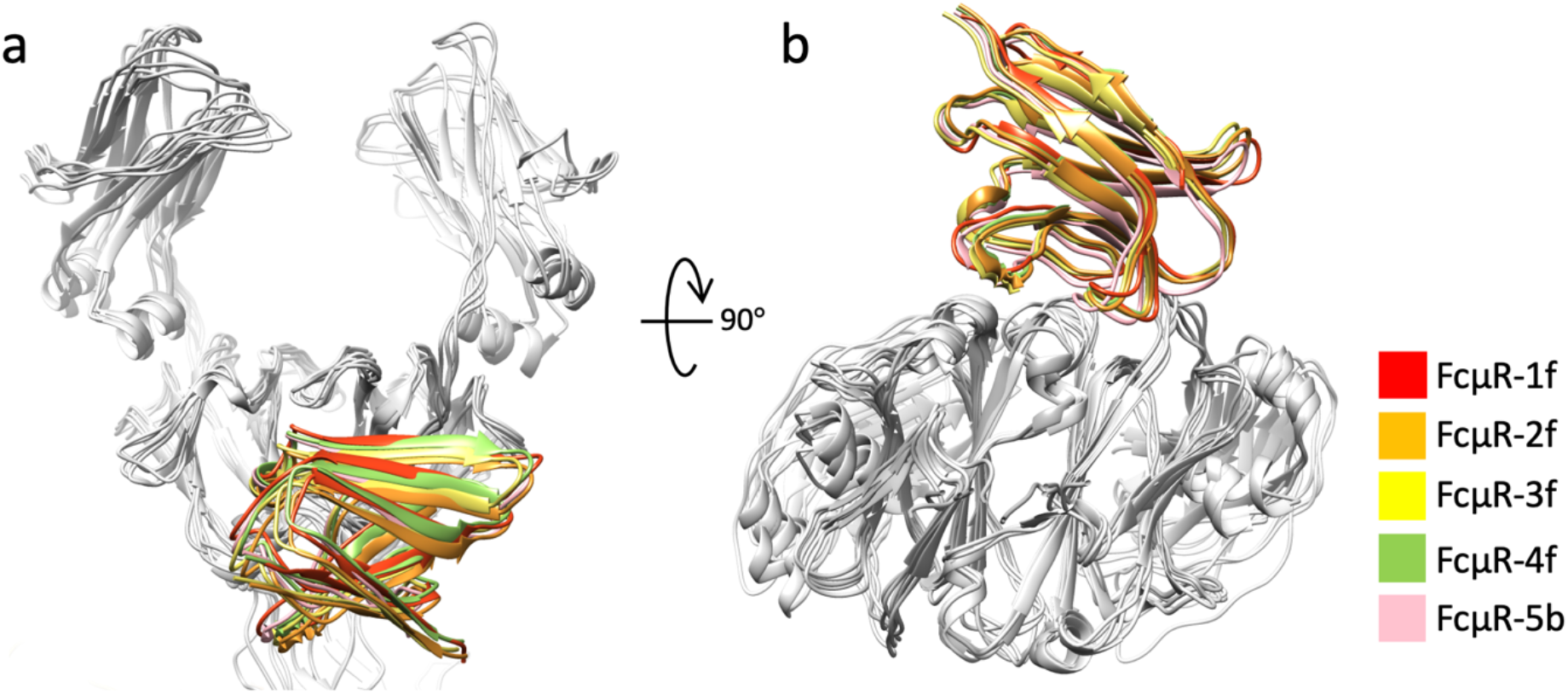
FcμR at different positions share similar positions relative to the IgM subunits. (a-b) Superposition of subunit Fcμ1 to Fcμ5 aligned at Fcμ chains showing the same binding position shared among the subunits. Taking FcμR-3f as reference, the backbone root mean squared deviation (RMSD) of FcμR-1f, FcμR-2f, FcμR-4f, and FcμR-5b is 2.6 Å, 1.4 Å, 1.6 Å, and 1.7 Å. FcμR-1f: FcμR bound at the front of Fcμ1 subunit. FcμR-2f: FcμR bound at the front of Fcμ2 subunit. FcμR-3f: FcμR bound at the front of Fcμ3 subunit. FcμR-4f: FcμR bound at the front of Fcμ4 subunit. FcμR-5b FcμR bound at the back of Fcμ5 subunit.

**Extended Data Fig. 4.**
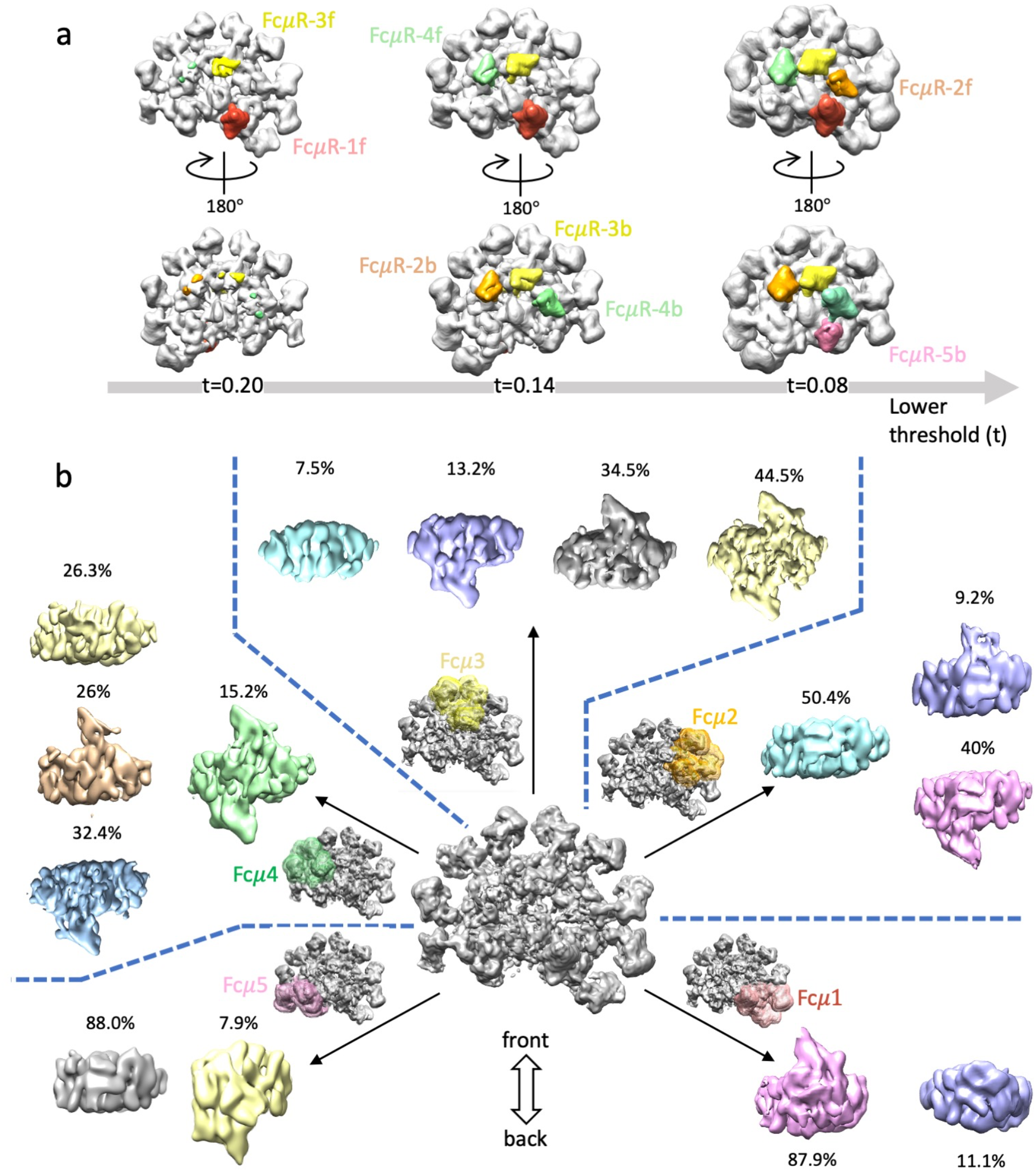
Different occupancies of FcμR among eight binding sites. (a) Gaussian-filtered (2 Å width) map (the non-uniform refined map in Extended Data Fig. 2a, before postprocessing) at three threshold levels showing the sequential appearance of FcμR densities from high to low occupancies. (b) Focused 3D classification at all subunits Fcμ1 to Fcμ5 for quantification of FcμR occupancy at each subunit. 3D classes of each subunit showing different FcμR binding states (at front, back, both or neither). Masks for particle subtraction at individual subunits are shown with different colours.

**Extended Data Fig. 5.**
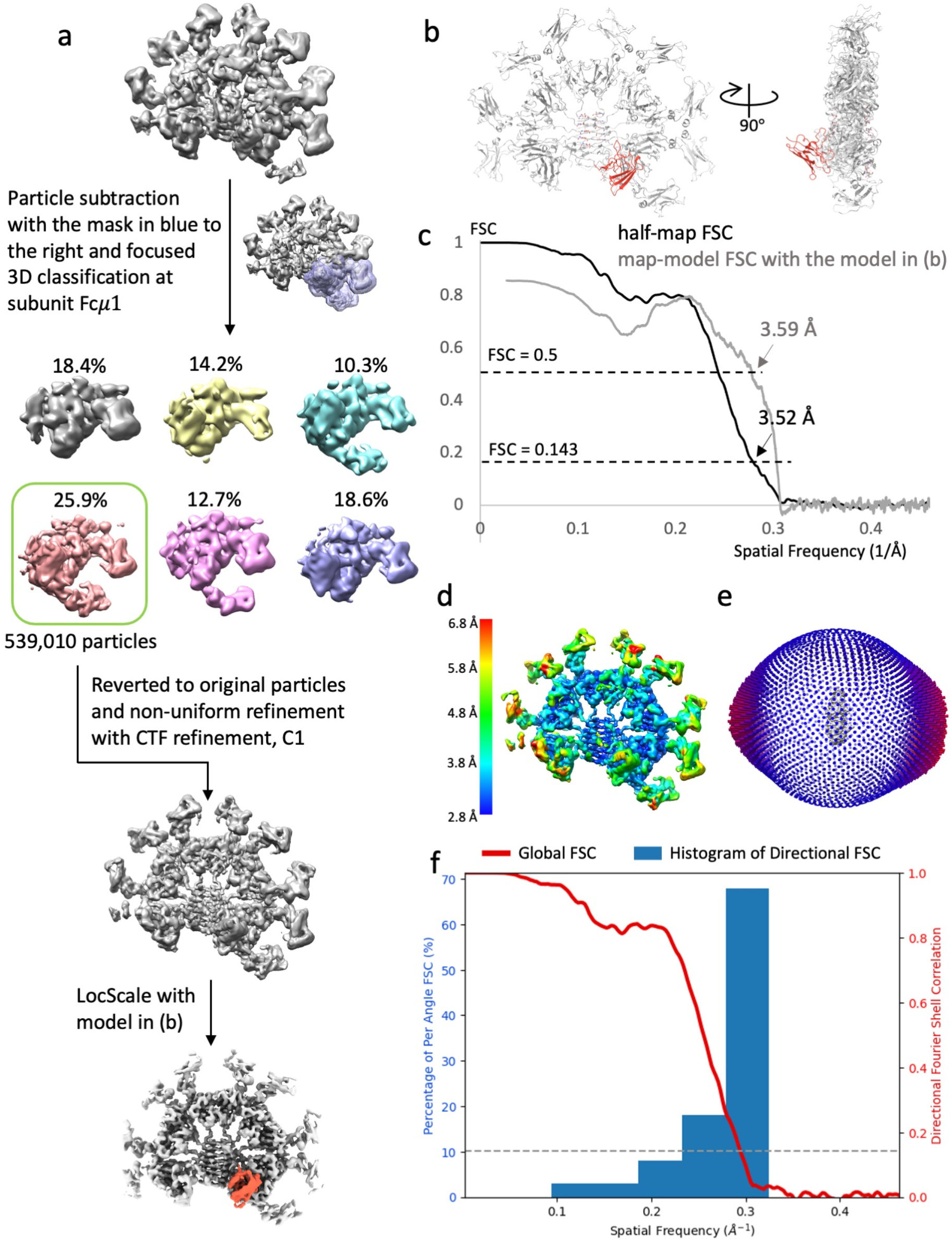
Cryo-EM structure of FcμR/IgM-Fc complex focused on subunit Fcμ1. (a) Particle subset selection by focused 3D classification at subunit Fcμ1 and map refinement. (b) Front and side view of the complex model. IgM in grey, FcμR in dark blue. (c) Gold standard Fourier shell correlation (FSC) with 3.5 Å resolution at 0.143 cut-off and map-model FSC plot showing 3.6 Å resolution at 0.5 cut-off calculated in Phenix. (d) Local resolution of the refined map calculated in Cryosparc. (e) Eulerian angle distribution of the particles in the non-uniform refinement. (f) 3DFSC histogram of the refined map calculated in Cryosparc of the refined map showing anisotropy between 3.2 Å -7.6 Å.

**Extended Data Fig. 6.**
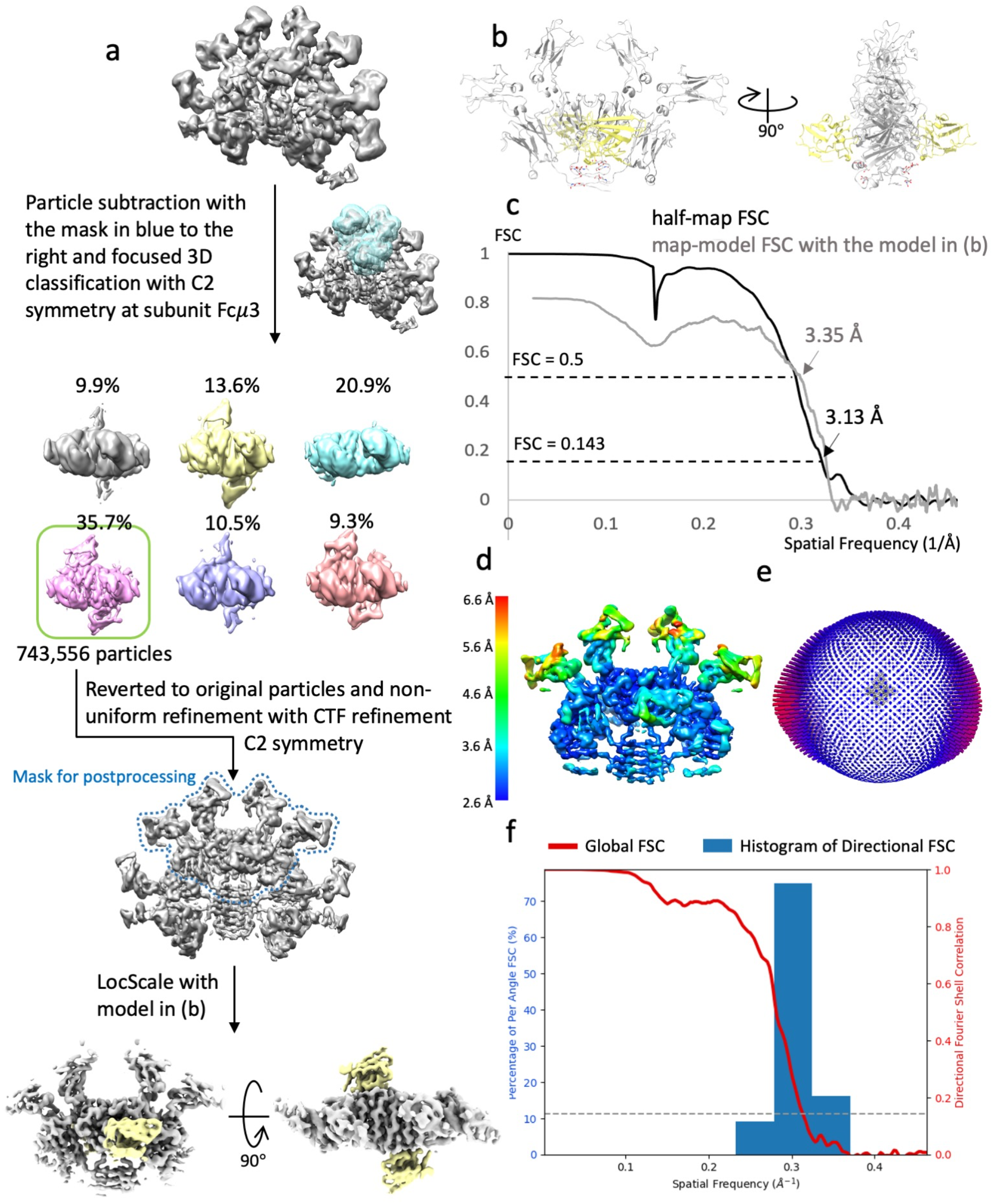
Cryo-EM structure of FcμR/IgM-Fc complex focused on subunit Fcμ3. (a) Particle subset selection by focused 3D classification at subunit Fcμ3 and map refinement. (b) Front and side view of the complex model. IgM in grey, FcμR in dark blue. (c) Gold standard Fourier shell correlation (FSC) with 3.1 Å resolution at 0.143 cut-off and map-model FSC plot showing 3.3 Å resolution at 0.5 cut-off calculated in Phenix. (d) Local resolution of the refined map calculated in Cryosparc. (e) Eulerian angle distribution of the particles in the non-uniform refinement. (f) 3DFSC histogram calculated in Cryosparc of the refined map showing angular resolution distribution from 3.0 Å - 3.8 Å.

**Extended Data Fig. 7.**
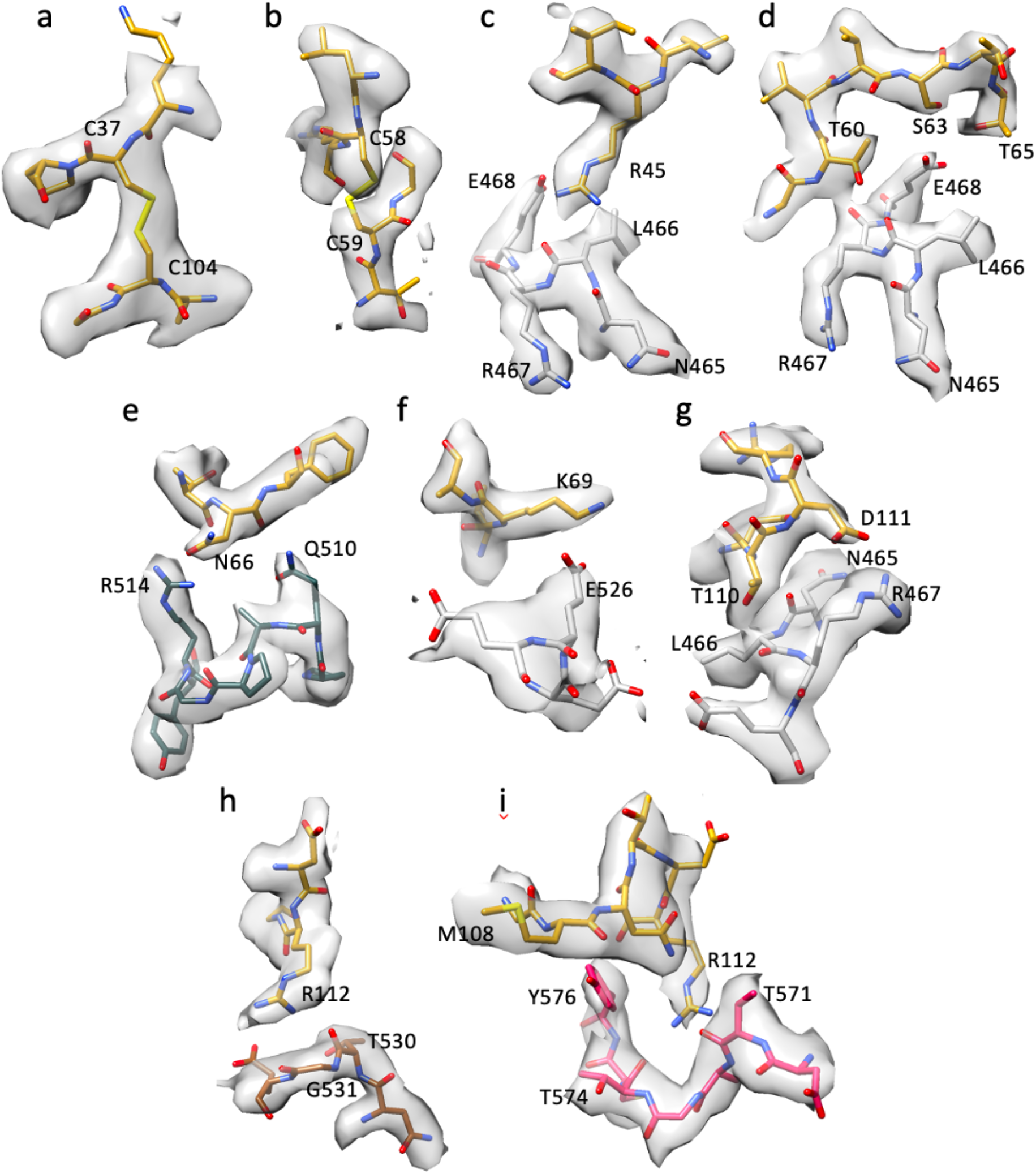
Density maps of key regions at FcμR and FcμR/IgM binding interface. (a-b) Two conserved disulfide bonds in FcμR. (c-g) Densities of the interacting residues on FcμR and Cμ4 domains, corresponding to the interaction shown in Fig. 2c. FcμR in dark yellow, Cμ4-B chain in light grey, and Cμ4-A chain in slate grey. (h) Densities of the residues in CDR3 regions of FcμR interacting with the neighbouring Cμ4 domain (in brown), corresponding to Fig. 2d. (i) Densities of the residues in CDR3 regions of FcμR interacting with the tailpiece of Fcμ5 chain (in pink), corresponding to Fig. 2e.

**Extended Data Fig. 8.**
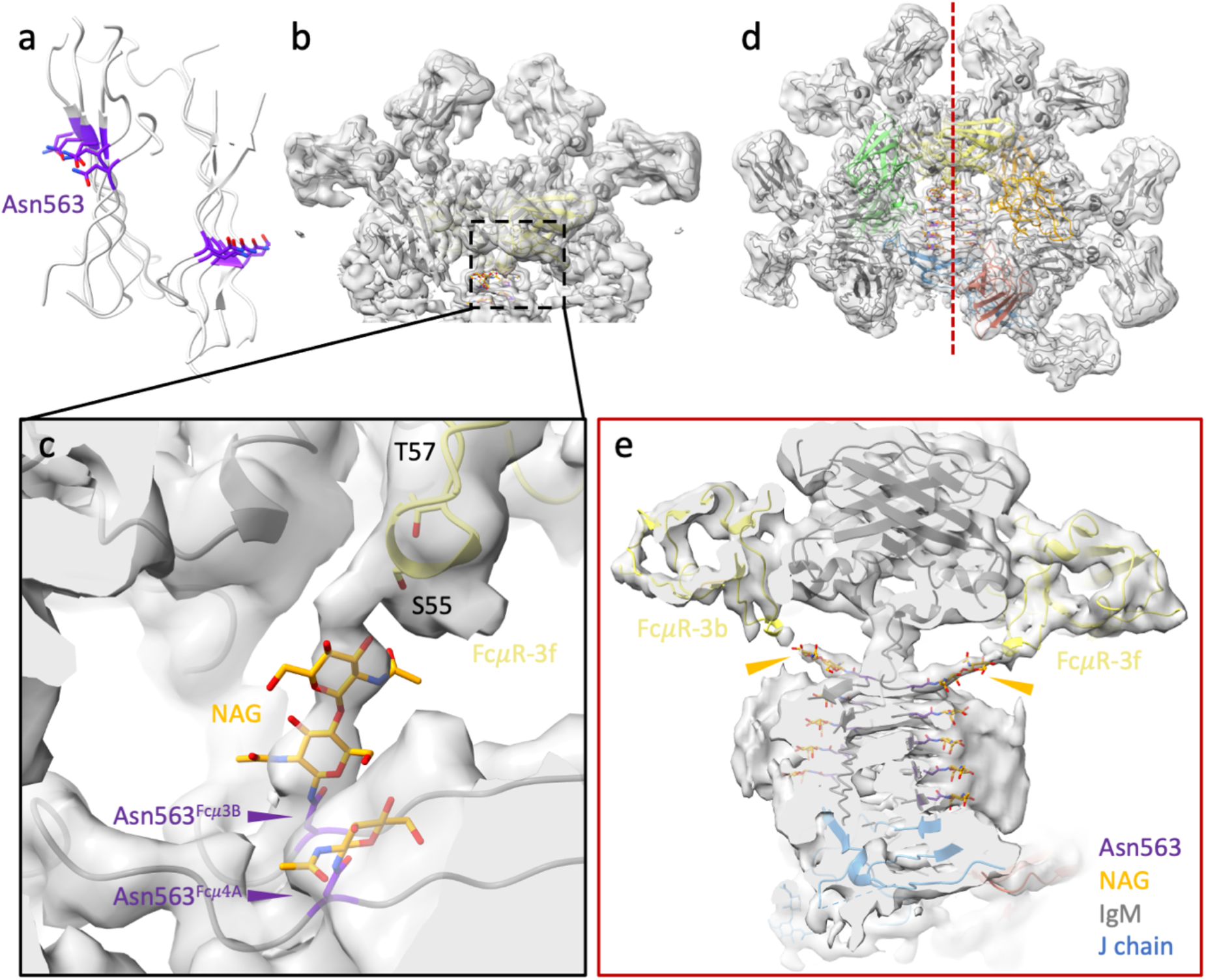
N-linked glycosylation at Asn563 contacting with FcμR at subunit Fcμ3. (a) The tailpiece assembly of IgM pentamer showing ten N-linked glycosylation sites (purple). (b) The map of subunit Fcμ3 (same map as Extended Data Fig. 6a, before postprocessing, map threshold=0.2). (c) Zoom-in view of the N-Acetylglucosamine (NAG) molecules (orange) linking from Asn563 (purple) at the tailpiece of Fcμ3B chain to FcμR-3f (yellow). (d) The overall map of FcμR/pIgM-Fc (same map as Extended Data Fig. 2a, before postprocessing, map threshold=0.2). (e) Cross-section of the map in (d) indicated by the red dotted line showing the densities of the two NAG chains (orange arrowheads) extending from Asn563 of the two Fcμ chains (Fcμ3A and Fcμ3B) to the two FcμR molecules at both sides.

**Extended Data Table 1.** Occupancy of FcμR on individual subunits of IgM pentamer based on 3D classification.

|  | Fcμ1 | Fcμ2 | Fcμ3 | Fcμ4 | Fcμ5 |
| --- | --- | --- | --- | --- | --- |
| at front | 87.9% | 9.2% | 79.0% | 41.2% | 0 |
| at back | 0 | 40% | 57.7% | 47.6% | 7.9% |
| at both | 0 | 0 | 44.5% | 15.2% | 0 |
| none | 11.1% | 50.4% | 7.5% | 26.3% | 88.0% |

**Extended Data Table 2.**
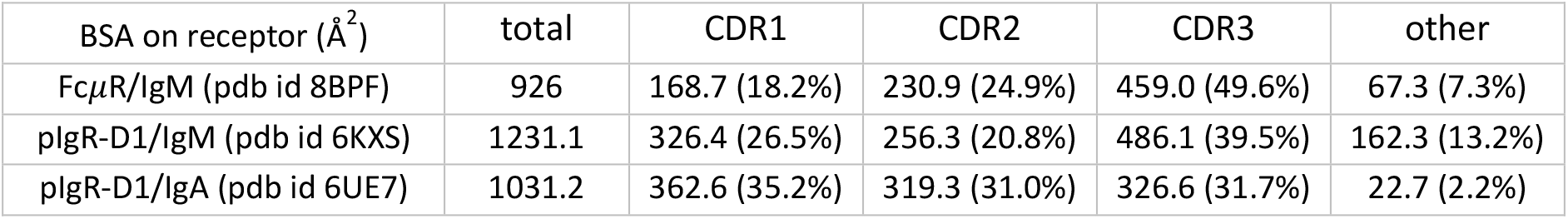
Buried surface areas between the individual CDR loops of the receptors and the immunoglobulin binding partner. The CDR regions are defined in the sequence alignment in Fig. 2f.

| BSA on receptor ($\text{\AA}^2$) | total | CDR1 | CDR2 | CDR3 | other |
| --- | --- | --- | --- | --- | --- |
| FcμR/IgM (pdb id 8BPF) | 926 | 168.7 (18.2%) | 230.9 (24.9%) | 459.0 (49.6%) | 67.3 (7.3%) |
| pIgR-D1/IgM (pdb id 6KXS) | 1231.1 | 326.4 (26.5%) | 256.3 (20.8%) | 486.1 (39.5%) | 162.3 (13.2%) |
| pIgR-D1/IgA (pdb id 6UE7) | 1031.2 | 362.6 (35.2%) | 319.3 (31.0%) | 326.6 (31.7%) | 22.7 (2.2%) |

**Extended Data Table 3.** Cryo-EM data collection, processing, and validation statistics.

|  |  |  |  |
| --- | --- | --- | --- |
|  | 8:1 FcμR/IgM-Fc (8BPE) | FcμR at subunit Fcμ1 (8BPF) | FcμR at subunit Fcμ3 (8BPG) |
| Data Acquisition |  |  |  |
| Voltage | 300 kV |  |  |
| Microscope | FEI Titan Krios |  |  |
| Camera | K2, counting |  |  |
| Calibrated magnification | 46,296 |  |  |
| Electron exposure | 50.6 e/Å² |  |  |
| Exposure rate | 5.06 e/Å²/s |  |  |
| Number of frames per movie | 40 |  |  |
| Energy filter slit width | 20 eV |  |  |
| Automation software | EPU |  |  |
| Stage tilt | 0° and 20° |  |  |
| Defocus range | -1.2 to -3.5 μm |  |  |
| Pixel size | 1.08 Å |  |  |
| Data processing |  |  |  |
| Data processing packages | Relion, CryoSPARC, CrYOLO, CCPEM, LocScale |  |  |
| Initial particle images | 5,923,734 |  |  |
| Symmetry imposed | C1 | C1 | C2 |
| Final particle images | 516,875 | 539,010 | 743,566 |
| Half-map FSC (0.143, masked, Å) | 3.63 | 3.52 | 3.13 |
| Map B factor (Å²) | 124 | 124 | 142 |
| Model Refinement |  |  |  |
| Initial model used | 6KXS, trRosetta predicted FcμR |  |  |
| Refinement/validation packages | Phenix, Coot, CCPEM, Privateer |  |  |
| Map-model FSC (0.5, masked, Å) | 3.87 | 3.59 | 3.35 |
| Map-model CC |  |  |  |
| CC_mask | 0.64 | 0.74 | 0.73 |
| CC_volume | 0.63 | 0.71 | 0.67 |
| CC_peaks | 0.56 | 0.63 | 0.54 |
| CC_box | 0.65 | 0.71 | 0.62 |
| Model composition |  |  |  |
| Non-hydrogen atoms | 24203 | 18594 | 8652 |
| Protein residues | 3104 | 2370 | 1106 |

| Ligands | NAG:14 | NAG:12 | NAG:6 |
| --- | --- | --- | --- |
| Validation |  |  |  |
| MolProbity score | 1.80 | 1.88 | 1.54 |
| Clashscore | 5.41 | 6.28 | 3.14 |
| Rotamer outliers (%) | 0 | 0 | 0 |
| Ramachandran outliers (%) | 0 | 0 | 0 |
| Cβ outliers (%) | 0 | 0 | 0 |
| Planarity outlier (%) | 0 | 0 | 0 |
| Chirality outlier (%) | 0 | 0 | 0 |
| Bond angle/length outlier (%) | 0 | 0 | 0 |

## References

1. Kubagawa, H. et al. Identity of the elusive IgM Fc receptor (FcμR) in humans. J. Exp. Med. 206, 2779–2793 (2009).

2. Akula, S., Mohammadamin, S. & Hellman, L. Fc receptors for immunoglobulins and their appearance during vertebrate evolution. PLoS One 9, e96903 (2014).

3. Kubagawa, H. et al. The long elusive IgM Fc receptor, FcμR. J. Clin. Immunol. 34, 35–45 (2014).

4. Li, Y. et al. Structural insights into immunoglobulin M. Science. 367, 1014–1017(2020).

5. Kumar, N., Arthur, C. P., Ciferri, C. & Matsumoto, M. L. Structure of the secretory immunoglobulin A core. Science. 367, 1008–1014 (2020).

6. Kumar, N., Arthur, C. P., Ciferri, C. & Matsumoto, M. L. Structure of the human secretory immunoglobulin M core. Structure 29, 1–8 (2021).

7. Wang, Y. et al. Structural insights into secretory immunoglobulin A and its interaction with a pneumococcal adhesin. Cell Res. 30, 602–609 (2020).

8. Shima, H. et al. Identification of TOSO/FAIM3 as an Fc receptor for IgM. Int. Immunol. 22, 149–156 (2009).

9. Lloyd, K. A., Wang, J., Urban, B. C., Czajkowsky, D. M. & Pleass, R. J. Glycan-independent binding and internalization of human IgM to FCMR, its cognate cellular receptor. Sci. Rep. 7, (2017).

10. Nyamboya, R. A., Sutton, B. J. & Calvert, R. A. Mapping of the binding site for FcμR in human IgM-Fc. Biochim. Biophys. Acta - Proteins Proteomics 1868, 140266 (2020).

11. Skopnik, C. M. et al. Identification of Amino Acid Residues in Human IgM Fc Receptor (FcμR) Critical for IgM Binding. Front. Immunol. 11, (2021).

12. Chen, Q., Menon, R., Calder, L. J., Tolar, P. & Rosenthal, P. B. Cryomicroscopy reveals the structural basis for a flexible hinge motion in the immunoglobulin M pentamer. Nat. Commun. 13, (2022).

13. Ma, X. et al. Cryo-EM structures of two human B cell receptor isotypes. Science. 377, 880–885 (2022).

14. Su, Q. et al. Cryo-EM structure of the human IgM B cell receptor. Science. 377, 875–880 (2022).

15. Dong, Y. et al. Structural principles of B-cell antigen receptor assembly. Nature doi: 10.1038/s41586-022-05412-7. Epub ahead of pri (2022).

16. Kubagawa, H. et al. Functional roles of the IgM Fc receptor in the immune system. Front. Immunol. 10, (2019).

17. Rochereau, N. et al. Essential role of TOSO/FAIM3 in intestinal IgM reverse transcytosis. Cell Rep. 37, (2021).

18. Ouchida, R. et al. FcμR Interacts and Cooperates with the B Cell Receptor To Promote B Cell Survival. J. Immunol. 194, 3096–3101 (2015).

19. Nguyen, T. T. T. et al. The IgM receptor FcμR limits tonic BCR signaling by regulating expression of the IgM BCR. Nat. Immunol. 18, 321–333 (2017).

20. Zivanov, J. et al. New tools for automated high-resolution cryo-EM structure determination in RELION-3. Elife 7, e42166 (2018).

21. Rohou, A. & Grigorieff, N. CTFFIND4: Fast and accurate defocus estimation from electron micrographs. J. Struct. Biol. 192, 216–221 (2015).

22. Punjani, A., Rubinstein, J. L., Fleet, D. J. & Brubaker, M. A. CryoSPARC: Algorithms for rapid unsupervised cryo-EM structure determination. Nat. Methods 14, 290–296 (2017).

23. Wagner, T. et al. SPHIRE-crYOLO is a fast and accurate fully automated particle picker for cryo-EM. Commun. Biol. 2, 218 (2019).

24. Du, Z. et al. The trRosetta server for fast and accurate protein structure prediction. Nat. Protoc. 16, 5634–5651 (2021).

25. Afonine, P. V. et al. New tools for the analysis and validation of cryo-EM maps and atomic models. Acta Crystallogr. Sect. D Struct. Biol. D74, 814–840 (2018).

26. Emsley, P., Lohkamp, B., Scott, W. G. & Cowtan, K. Features and development of Coot. Acta Crystallogr. Sect. D Biol. Crystallogr. 66, 486–501 (2010).

27. Jakobi, A. J., Wilmanns, M. & Sachse, C. Model-based local density sharpening of cryo-EM maps. Elife 6, (2017).

28. Burnley, T., Palmer, C. M. & Winn, M. Recent developments in the CCP-EM software suite. Acta Crystallogr. Sect. D Struct. Biol. 73, 469–477 (2017).

29. Agirre, J. et al. Privateer: Software for the conformational validation of carbohydrate structures. Nat. Struct. Mol. Biol. 22, 833–834 (2015).

30. Pettersen, E. F. et al. UCSF Chimera - A visualization system for exploratory research and analysis. J. Comput. Chem. 25, 1605–1612 (2004).

31. Goddard, T. D. et al. UCSF ChimeraX: Meeting modern challenges in visualization and analysis. Protein Sci. 27, 14–25 (2018).

